# Effect of egg production dynamics on the functional response of parasitoids

**DOI:** 10.1101/2023.03.22.533781

**Authors:** María Aguirre, Guillermo Logarzo, Serguei Triapitsyn, Hilda Diaz-Soltero, Stephen Hight, Octavio Bruzzone

## Abstract

Functional response describes the number of hosts attacked by a parasitoid in relation to host densities and plays an important role by connecting behavioral-level processes with community-level processes. Most functional response studies were carried out using simple experimental designs where the insects were confined to a plain and small arena with different host densities during a fixed period of time. With these designs, other factors that might affect the functional response of parasitoids were not analyzed, such as fecundity, age, and experience. We proposed a series of latent-variables Markovian models that comprised an integrated approach of functional response and egg production models to estimate the realized lifetime reproductive success of adult parasitoids. As a case study, we used the parasitoids *Anagyrus cachamai* and *A. lapachosus* (Hymenoptera: Encyrtidae), two candidate agents for neoclassical biocontrol of the Puerto Rican cactus pest mealybug, *Hypogeococcus* sp. (Hemiptera: Pseudococcidae). *Anagyrus cachamai* and *A. lapachosus* presented a type III functional response. However, the two parasitoids behaved differently: for *A. cachamai*, the number of parasitized hosts decreased with female age and depended on the number of mature eggs that were available for oviposition, whereas *A. lapachosus* host parasitism increased with female age and was modulated by its daily egg load and previous experience. The tested species were assessed according to their physiology and prior experience. We estimated the number of mature eggs after emergence, egg production on the first day, egg production rate, proportion of eggs resorbed, egg resorption threshold, and egg storage capacity. The methodology presented may have large applicability in pest control, invasive species management, and conservation biology, as it has the potential to increase our understanding of the reproductive biology of a wide variety of species, ultimately leading to improved management strategies.

## Introduction

Functional response is one of the most commonly used mathematical frameworks to describe and estimate the number of hosts attacked by a parasitoid in relation to host densities [1,2, allowing the connection of behavioral-level processes with community-level processes. Identifying a parasitoid’s response to changes in host-density is central to any description of parasitism because the number of hosts attacked determines development, reproduction, and survival of the parasitoid [3]. Applications of this mathematical framework are found in studies on pest control, invasive species management, and conservation biology [4].

Despite the abundance of functional response models [5], the most frequently used are Holling’s type II and type III [1], describing a hyperbolic saturating curve and a sigmoid curve, respectively. Development of both functional response models requires only two parameters: *a*, the attack rate, and *H*, handling time. The attack rate represents the efficiency of a parasitoid in locating hosts via different “areas of discovery” measured in various habitat complexities or experimental arena sizes, according to Rogers [6], or simply, efficiency in locating hosts, measured as the proportion of hosts found per time unit according to Holling [1]. Handling time, *H*, is the time that a parasitoid spends manipulating its hosts. According to Holling [1], its inverse is the asymptotic value of the functional response curve. Type III sigmoidal functional response curves represent situations where the parasitoid is assumed to switch between two or more host species as a result of host availability or learning [1,7, but can also represent an improved efficiency by learning even when only one host is present [8].

In simple terms, attack rate describes the space that a parasitoid seeks per unit of time, while handling time is associated with host processing. Functional response models assume continuous foraging by individuals, along with stationary behavioral and physiological processes, when actually a plethora of biological processes are included under these two parameters [9,10. Most parasitoid functional response studies are carried out using experimental designs where the insects are confined to a small arena with different host densities during a fixed time period ranging from 1 to 48 hours [11–16]. These experimental designs ignore factors related to parasitoid behavior that affect functional response, such as fecundity, age, and experience of the wasp [17–19]. Varone et al. [20], studying functional response throughout the entire female lifetime of the larval parasitoid *Campoletis grioti* Blanchard (Hymenoptera: Ichneumonidae), found that attack rate and handling time of the parasitoid varied throughout the female’s lifespan, determined by the daily load of mature eggs.

Parasitoid wasps exhibit a wide spectrum of reproductive strategies that lead to variation in egg production dynamics. Egg load varies throughout the female’s lifespan, responding to both individual physiological and environmental factors [21–25]. In this context, the number of eggs that a female lays during her lifetime is determined by the number of hosts that the female encounters, the number of mature eggs over the female’s life span, and the behavior affecting the oviposition rate [26]. Since egg production is costly, selection should favor production strategies in which parasitoid females do not die before exhausting their egg complement (time limitation) or run out of eggs before all available hosts are used (egg limitation) [27]. Understanding ovarian dynamics is particularly relevant to describing the parasitoids’ foraging behavior because the physiological status of the ovaries may determine, for example, the duration of the pre-reproductive period and the rate of oviposition.

Egg limitation is mediated by oviposition and ovarian production, which in turn is regulated by two processes: egg maturation and egg resorption [28]. Parasitoid longevity influences the extent to which parasitoid becomes time-limited. To maximize longevity in the field, many parasitoids require a carbohydrate source such as nectar, hemipteran honeydew, or they might feed directly on their hosts [29,30. However, the variability in nutrient income caused by the use of external stochastic sources of nutrients can entail great risks of starvation. Egg resorption acts as an insurance against stochasticity, but it is considered as a “last–resort” strategy given the relatively low energy content of an egg [31,32. What distinguishes resorption from other sources of nutrients is its controllable nature, since the reserves contained in the eggs are made readily available to the female when they are most needed.

For over 80 years, researchers have proposed different models to determine if the realized lifetime reproductive success of adult female parasitoids was limited by the finite amount of time available to locate hosts that serve as oviposition sites [33–37], or by the finite supply of mature eggs [38,39. Rosenheim [40], using models to explore how stochasticity influences the evolution of egg limitation in insects, found that both egg and time limitations are fundamental in shaping insect reproductive behavior and population dynamics. These results underscore the importance of developing models of insect reproduction and population dynamics that incorporate the constraints imposed by both egg and time limitation, rather than just one constraint or the other.

In this study, we proposed a series of latent-variables Markovian models that comprised an integrated approach of functional response and egg production models to estimate the realized lifetime reproductive success of adult parasitoids. As a case study, we used the parasitoids *Anagyrus cachamai* Triapitsyn, Logarzo & Aguirre and *A. lapachosus* Triapitsyn, Aguirre & Logarzo (Hymenoptera: Encyrtidae), promising candidates for the neoclassical biological control of the mealybug *Hypogeococcus* sp. (Hemiptera: Pseudococcidae), a pest of native cacti in Puerto Rico [41–43]. Both parasitoid species are synovigenic wasps that do not engage in host feeding, but each presents differences in their reproductive biology (egg load at birth, ovigeny index and sex ratio) [42]. Each parasitoid species was evaluated employing a dynamic variant of the most common functional response models (e.g., [1,44), which included population parameters related to the parasitoids’ fecundity (egg resorption and daily egg load, limited by egg load capacity and daily egg production), and the age of the female parasitoid. Unlike the techniques used in previous classical methods, we worked on the entire parasitoid lifetime and incorporated physiological processes related to egg load into our evaluation of parasitism efficiency. This technique will provide a more accurate estimation between the two tested parasitoid species according to their physiology and prior experiences.

## Materials and Methods

The studies were conducted at the Fundación para el Estudio de Especies Invasivas (FuEDEI), located in Hurlingham, Buenos Aires, Argentina, between January 2014 and December 2016. All experiments and insect rearing were carried out in environmental-controlled chambers (25 ± 1°C, 16:8 L:D, 60–80% RH). All observations were conducted under a dissecting microscope at 40X.

### Parasitoid rearing

Laboratory experiments were conducted with colonies of *A. cachamai* and *A. lapachosus* reared at FuEDEI since 2014 following the methodology described in Aguirre et al. [42]. Each primary parasitoid species was reared on first instar nymphs of *Hypogeococcus* sp. “*Cactaceae host-clade*” [45], a congener but a different species from the mealybug pest of cacti in Puerto Rico [43]. Pure mealybug colonies were reared without parasitoids on clean potted plants of *Cleistocactus baumannii* (Lem.) Lem. (Cactaceae).

Colonies of *A. cachamai* and *A. lapachosus* were reared in separate rooms. Four mated females of each wasp species were placed in a plastic cage (2 L) with a 6 cm diameter hole cut in the lid and covered with polyester gauze for ventilation. The cage contained a piece of *C. baumannii* (20-25 cm long) infested with about 100 nymphs of *Hypogeococcus* sp. “*Cactaceae host-clade*”. After 72 hours, the four female parasitoids were removed from the plastic cage, and the parasitoid-exposed nymphs were monitored every three days. After the first parasitoid pupa was detected, monitoring was conducted daily, and all parasitoid pupae found were transferred to a Petri dish (1.5 cm high x 5.5 cm diameter) covered with plastic food wrap to keep wasps from escaping after emergence. Using this process, the wasps’ age, feeding conditions, and mating were controlled. As the parasitoids emerged, they were placed in a new Petri dish of equal dimensions with a squashed drop of honey on the bottom, and covered with clear plastic food wrap, to be used either for rearing or experimental purposes. The age of the female parasitoids for the experiments was 24 hours old; they were fed, mated, and had no previous oviposition experience. Throughout the paper, the mention of *Hypogeococcus* sp. nymphs exposed to female parasitoids refers to first instar nymphs of *Hypogeococcus* sp. “*Cactaceae host-clade*” on 20-25 cm long pieces of *C. baumannii*.

### Functional response experiments

To estimate the functional response of the parasitoids *A. cachamai* and *A. lapachosus*, a constant daily density of non-parasitized *Hypogeococcus* sp. nymphs was exposed to a female parasitoid once she was 24 hours old and until her death [20]. The daily number of parasitized nymphs was estimated for each female by recording the number of emerged parasitoids relative to the number of nymphs offered. Six nymph densities were evaluated: 10, 20, 40, 60, 80 and 110, with 5 replications per density, and a maximum error of 10% in the daily number of nymphs per density offered. The densities selected in this study were based on the results of a pilot test, where densities of 80 and 110 nymphs produced a plateau in the curve of the number of nymphs attacked as a function of the host density offered.

The experiments were conducted in vented plastic cages similar to the one described above for parasitoid rearing. To ensure that the daily number of non-parasitized nymphs available to each wasp was constant, the cactus piece with the nymphs exposed to the wasp was removed every 24 hours from the experimental arena and replaced by another piece of cactus with an equal number of nymphs not previously exposed to a parasitoid. Each cactus piece with the parasitoid-exposed nymphs was held individually in a similar plastic cage. All exposed nymphs were checked every three days and the number of emerged parasitoids counted until all non-parasitized nymphs completed their development and all wasps had emerged from parasitized nymphs.

## Data analyses

### Description of models

The outcome of the functional response experiments was analyzed with a series of latent-variables Markovian models that comprised an integrated approach of functional response and egg production models. Each model was summarized in a single equation that integrated two modules, one represented a functional response equation and another an equation of egg production:

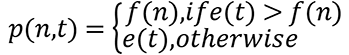

here *f*(*n*) represents the functional response equation, *e*(*t*) the egg production equation, and *p*(*n*,*t*) the model that describes the number of eggs laid by a wasp. If the number of eggs that a female has available is greater than the number of eggs that can be oviposited according to the functional response module, then the number of hosts attacked is predicted by the functional response equation. If the females’ egg load is less than the number of hosts available, she simply lays all the eggs she has. Using this basic structure, the proposed models are the combination of one of the six functional response equations of the functional response module with one of the eight egg production equations of the egg production module. This resulted in a matrix of 48 possible models, all of which were tested. Both modules are briefly described below.

#### Functional response module

The six equations tested in the functional response module were based on type I, type II, and two type III generalized functional responses [1,8,44,46–48]. For each of the type III generalized functional response equations, an additional version where the female gains experience in the course of her life when interacting with the host was also proposed. See the appendix section in the supporting information for details on the six equations tested (S1 File, Functional response module).

#### Egg production module

Eight egg production equations were proposed and tested. The simplest equation considers that the female has unlimited egg production. The equation with the next level of complexity assumes that the female is strictly pro-ovigenic and therefore all of its oocytes are mature upon emergence. The remaining six equations describe the behavior of synovigenic females, meaning, females that emerge with few or no mature eggs but continue to mature eggs throughout their lifetime. The most complex synovigenic-based model includes parameters related to egg resorption and daily egg load, limited by egg load capacity and daily egg production. The eight egg production equations proposed for testing within the egg production module are presented in the appendix section in supporting information (S1 File, Egg production module).

### Model fitting and selection

We used a fully Bayesian approach and the Metropolis-Hastings algorithm [49,50 in order to select the best explaining models (out of the 48 models proposed) and to calculate their parameters. Traditionally, statistical analysis of the functional response experiments is carried out by selecting the functional response model by a logistic regression thereby reducing the problem of differentiating between a hyperbolic curve (type II functional response) and a sigmoid curve (type III functional response). The use of a non-linear regression in a frequentist framework is then recommended to estimate the parameters of the curve [51]. Since this approach is not appropriate for selecting several models that compete with one another, Johnson and Omland [52] proposed the use of the Bayesian system. In this work, the selection of models and the estimation of parameter distribution were conducted in a Bayesian framework. The results of the analysis enabled us to infer with which models and parameters it is possible to explain the results of the laboratory experiments and their statistical distributions.

The Deviation Information Criterion (DIC) index was used as the decision rule for model selection [53]. The models that presented lower DIC were selected according to Gelman et al. [54], as a balance of the explanatory power (in terms of the likelihood function) and complexity (in terms of number of parameters). It is necessary to obtain DIC values that have a difference greater than 5 among the different models in order to select one model over other models. If DIC values among the different models are not greater than 5, then model averaging is required following Burnham and Anderson [55]. A total of 200,000 iterations were used to fit the models; the first 100,000 were discarded as a “burn-in” for model selection, and the remaining iterations were used to calculate the parameters of each model and the information indexes. The a *priori* distribution of the parameters of the functional response curves were normal distributions with a mean of 0 and a variance equal to 100, or uniform distributions defined between the minimum and the maximum value that each parameter can obtain. In some parameters, such as handling time (*H*), attack rate of the female when she emerges (*b*), the number of mature eggs when a female emerges (*h*_0_), egg resorption threshold (*u*), and egg storage capacity (*C*), the values were restricted to be positive, since negative values would not make biological sense. Since the variables obtained (number of parasitized hosts) are discrete and bounded, the binomial likelihood function was used [54]. Finally, fitness of each of the selected models to the data of the experiments was calculated by using the generalized coefficient of determination (GCD) for binary data, according to Cox and Snell [56] and Magee [57].

The analyses were carried out by using a Parasitoid-Egg model version 0.02 [58] for the parasitoid model, and PyMC version 2.3.7 for Monte Carlo methods [59] for parameter calculation and fitting.

## Results

From the 48 models proposed to explain the pattern observed in the functional response experiments conducted with the parasitoids *A. cachamai* and *A. lapachosus* (Figs 1 and 2), four models were selected for each species (S1 Table). In both encyrtid species, the difference observed in the DIC values of the selected models was lower than 5; consequently, it was not possible to select a unique model for each species. *Anagyrus cachamai* and *A. lapachosus* females presented a type III functional response, but they used different strategies to exploit their host, *Hypogeococcus* sp. nymphs. On the other hand, although both species are synovigenic, females showed differences in their pattern of egg production though their lifetime.

**Fig 1.**
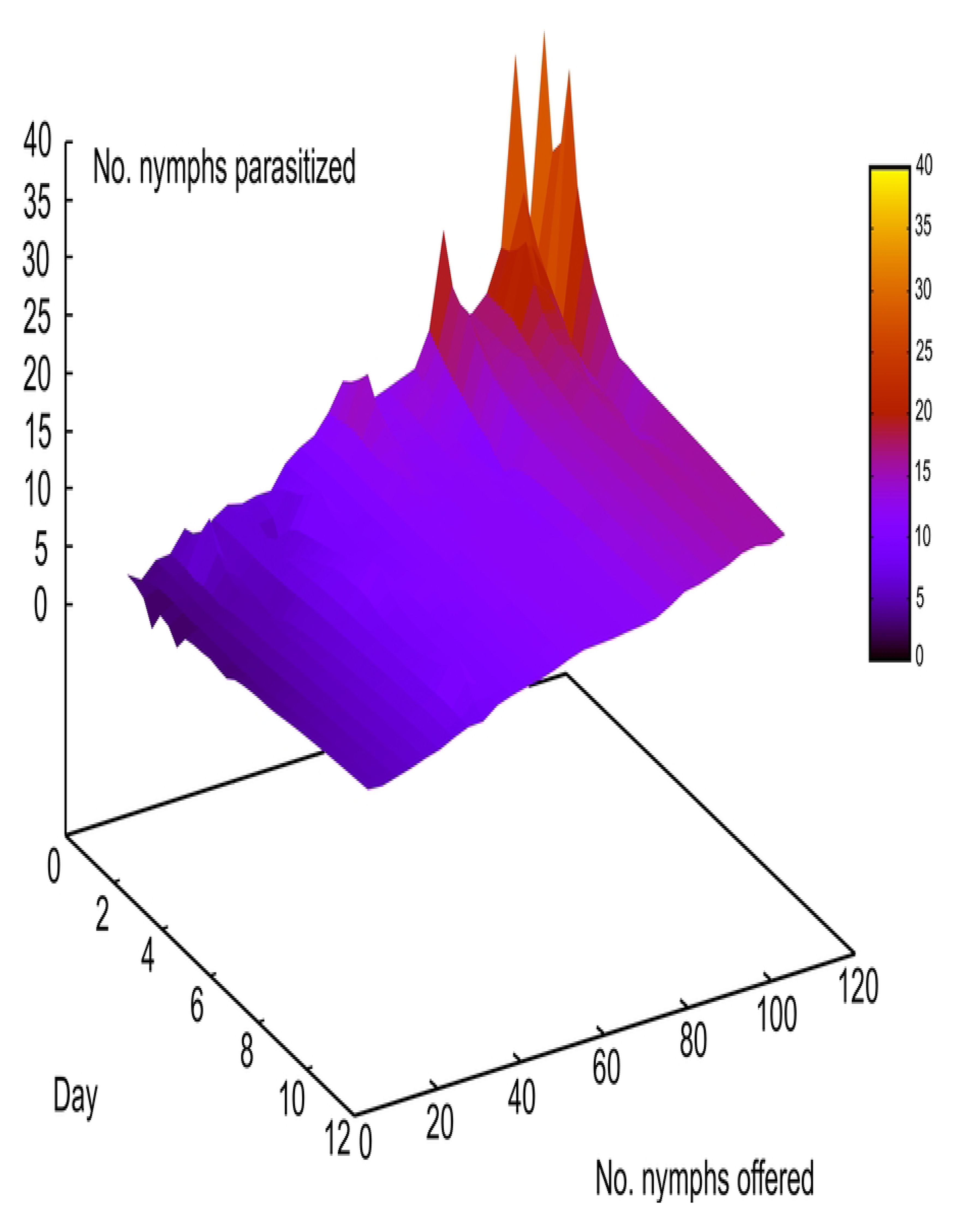
Observed functional response of *Anagyrus cachamai* females at different ages (days) of their lifespan (1-11 days) represented as the total number of nymphs parasitized relative to the total number of *Hypogeococcus* sp. nymphs offered. Six nymph densities were evaluated: 10, 20, 40, 60, 80 and 110, with 5 replications per density, and a maximum error of 10% in the number of nymphs per density offered.

**Fig 2.**
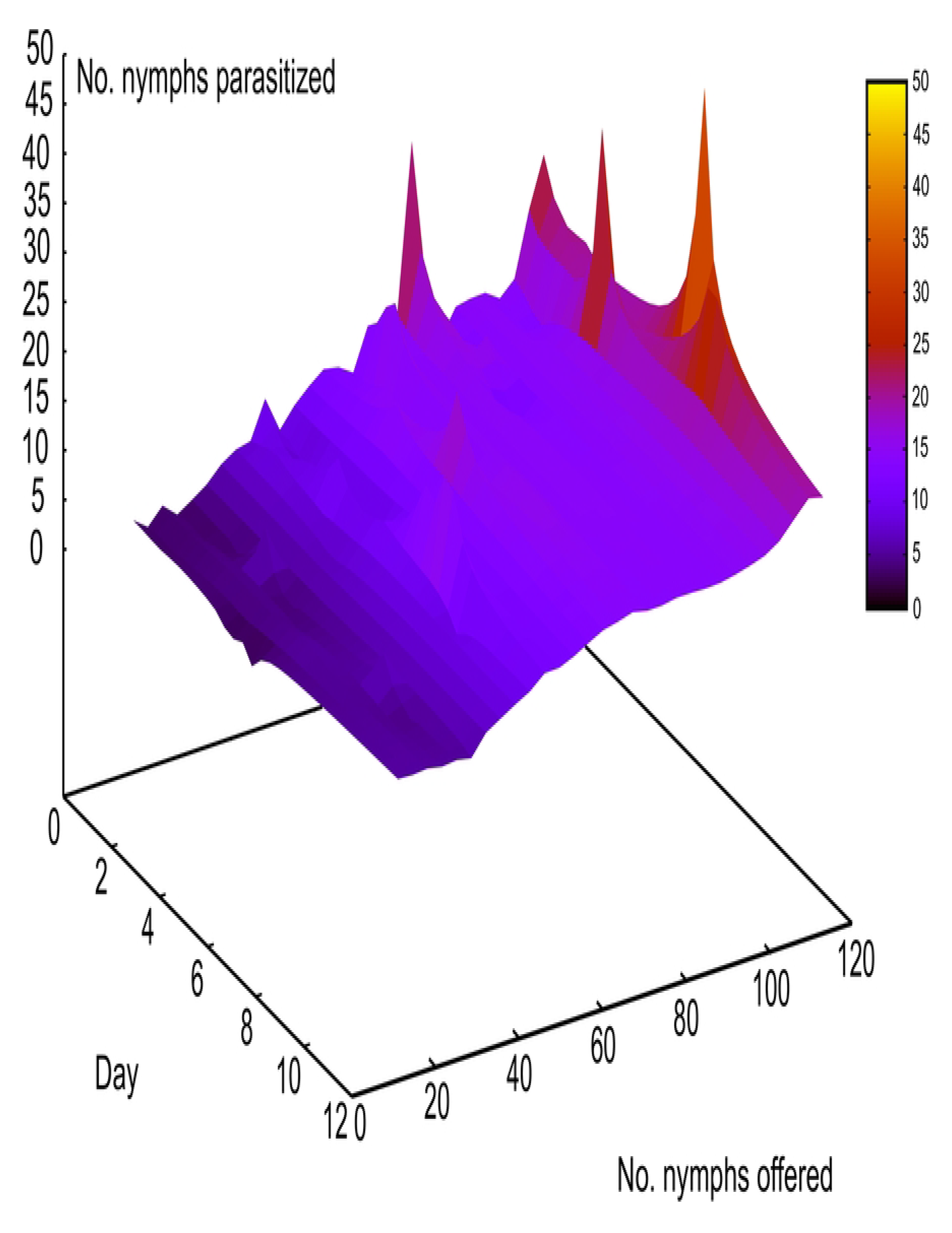
Observed functional response of *Anagyrus lapachosus* females at different ages (days) of their lifespan (1-12 days) represented as the total number of nymphs parasitized relative to the total number of *Hypogeococcus* sp. nymphs offered. Six nymph densities were evaluated: 10, 20, 40, 60, 80 and 110, with 5 replications per density, and a maximum error of 10% in the number of nymphs per density offered.

The following models were selected for the parasitoid *A. cachamai*: *C*5 (DIC = 1933.43), *E*5 (DIC = 1932.39), *C*7 (DIC = 1933.38) and *E*7 (DIC = 1935.21) (S1 Table); and their explanatory power in terms of GCD was GCD_*C*5_ = 0.85, GCD_*E*5_ = 0.85, GCD_*C*7_ = 0.85, and GCD_*E*7_ = 0.86, respectively. The four integrated models selected comprised between 8-9 parameters. When the components of the four selected models were analyzed in terms of functional response, two type III functional response equations were selected (Table 1, Fig 3A and 3B). In those models where equation *C* of the functional response module was selected, the attack rate increased linearly with the number of hosts offered. Emerged female attack rate was 0.073 ± 0.015 days^-1^ (*b*), the attack rate change was 0.003 ± 0.000 days^-1^(*a*), and the handling time was 0.005 ± 0.001 days (*H*). In the models where the equation *E* of the functional response module was selected, the attack rate changed with host densities as *an*^*s*^, where *a* = 0.018 ± 0.005 days^-1^, *s* = 1.676 ± 0.063 and *H* = 0.005 ± 0.001 days. In reference to the egg production module, two equations were selected; equation 5 and equation 7. *Anagyrus cachamai* females emerged with 56 ± 2 mature eggs (*e*), and lived on average 4 ± 2 days (range 2-11 days). During the first day of life, a female produced 8 ± 1 eggs (*h*_0_), and the daily egg production rate was 0.972 ± 0.036 (*g*). Eggs that were not used on day *t*, were resorbed on day *t* + 1, providing the number of accumulated eggs from one day to another was greater than 19 ± 1 eggs (*u*). When resorption existed, the proportion of eggs resorbed was 0.677 ± 0.045. In equation 7, females also presented an egg storage capacity of 58 ± 2 eggs (*C*). The four integrated models selected are detailed in S2 Table and represented in S1-S4 Figs.

**Fig 3.**
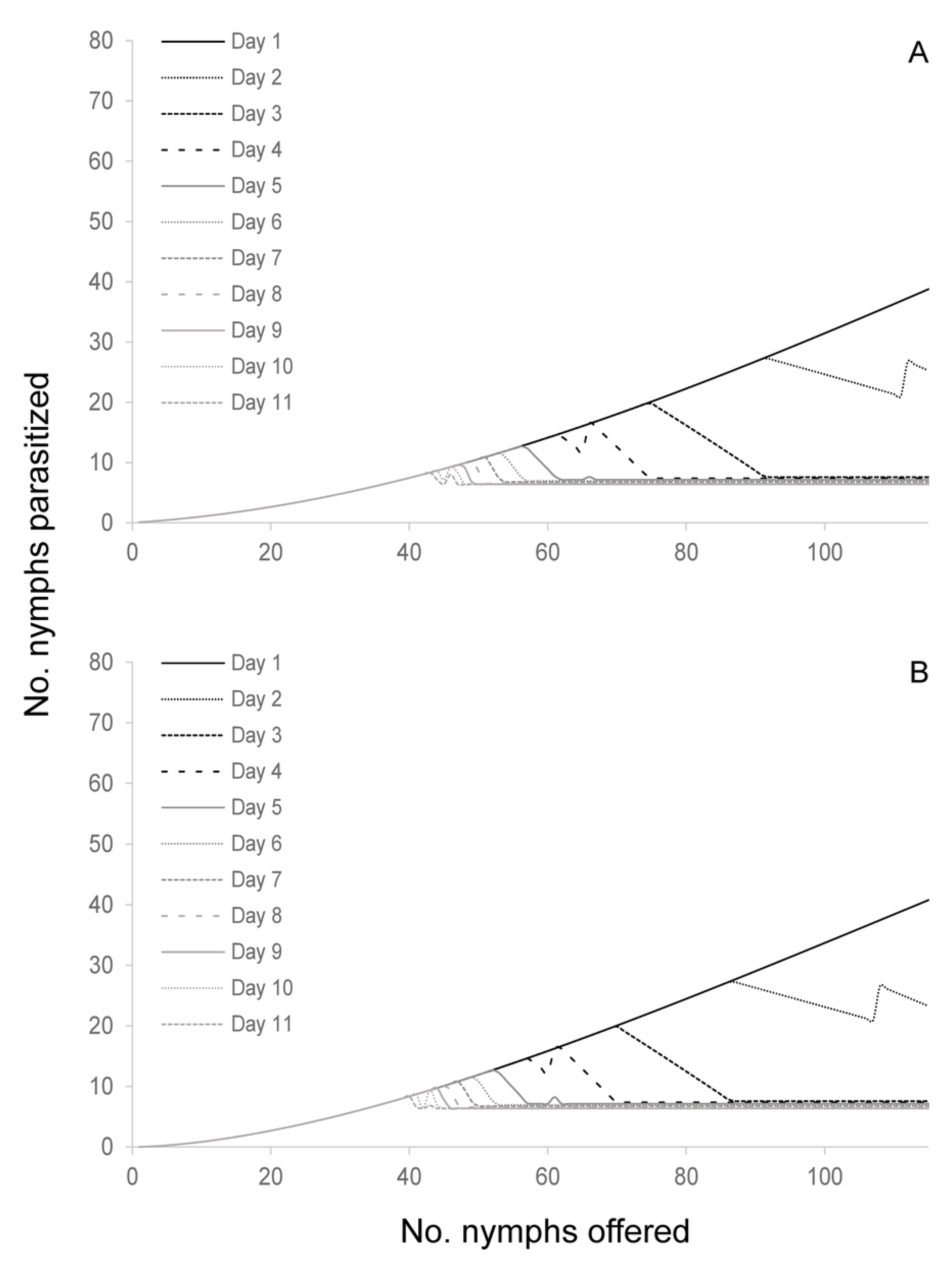
Estimated type III functional response of *Anagyrus cachamai* females at different ages (days) of their lifespan (1-11 days) considering limited egg production. (A) Equation *C* of functional response module; the attack rate increases linearly with the number of hosts available [46,47; (B) Equation *E* of functional response module; the attack rate changes with host densities as *an*^*s*^[44].

**Table 1.** A posteriori mean ± standard deviation of the species-specific parameters of the selected models for two rasitoids, Anagyrus cachamai and A. lapachosus, attacking Hypogeococcus sp. The values presented are the result of eighing the models selected for *A. cachamai* and *A. lapachosus* females (see the supporting information for details on the models lected: S1-S3 Tables).

|  | Functional response module parameters |  |  |  |  |  |  |  |  |
| --- | --- | --- | --- | --- | --- | --- | --- | --- | --- |
|  | Without female experience |  |  |  |  |  | With female experience |  |  |
| | FR III (attack rate increased linearly with the available host number [46,47]) | | | FR III (the attack rate changed with host densities as $\alpha n^s$ [44]) | | | FR III (attack rate increased linearly with the available host number [46,47]) | | |
| | Attack rate after emergence at $n=0$ ($b$) | Attack rate change ($a$) | Handling time ($H$) | Attack rate change ($a$) | Exponent $s$ ($1+q$) | Handling time ($H$) | Attack rate after emergence at $n=0$ ($b$) | Attack rate change ($a$) | Handling time ($H$) |
| *A. cachamai* | 0.073±0.015 $d^{-1}$ | 0.003±0.000 $d^{-1}$ | 0.005±0.001 $d$ | 0.018±0.005 $d^{-1}$ | 1.676±0.063 $d^{-1}$ | 0.005±0.001 $d$ | - | - | - |
| *A. lapachosus* | - | - | - | - | - | - | 0.109±0.010 $d^{-1}$ | 0.001±0.000 $d^{-1}$ | 0.004±0.001 $d$ |
|  | Egg production module parameters |  |  |  |  |  |  |  |  |
| | No. mature eggs after emergence ($e$) | Egg prod. on the first day ($h_0$) | Egg production rate ($g$) | Proportion of eggs resorbed ($r$) | Egg resorption threshold ($u$) | Egg storage capacity ($C$) | | | |
| *A. cachamai* | 56±2 | 8±1 | 0.972±0.036 | 0.677±0.045 | 19±1 | 58±2 | | | |
| *A. lapachosus* | 57±2 | 11±1 | 1.330±0.024 | 0.935±0.028 | 15±1 | 59±2 | | | |

Parameters reported for the egg production term are averaged considering all the iterations with the functional response type III thout female experience in the case of the species *A. cachamai* or with experience for *A. lapachosus*. On the other hand, the nctional response parameters were not mixed because the values and their behavior were slightly different depending on the kind model selected. Physical units of the calculated parameters: *d* is days, parameters without units are dimensionless.

The models selected for the species *A. lapachosus* were *D*4 (DIC = 2974.66), *D*5 (DIC = 2973.60), *D*6 (DIC = 2976.63), and *D*7 (DIC = 2975.19); and their explanatory power in terms of GCD was GCD_*D*4_ = 0.86, GCD_*D*5_ = 0.86, GCD_*D*6_ = 0.86, and GCD_*D*7_ = 0.86, respectively. The number of parameters of the four models selected ranged between 7-9. For the functional response module, equation *D* was selected which assumes that the female has a type III functional response and that the efficiency increases linearly with the number of hosts offered during the females’ life. This meant a daily increase in the attack rate of *A. lapachosus* females (Table 1, Fig 4). Emerging female attack rate was 0.109 ± 0.010 days^-1^ (*b*), the daily attack rate change was 0.001 ± 0.000 days^-1^ (*a*), and the handling time was 0.004 ± 0.001 days (*H*). Regarding the egg production module, four egg production equations were selected: 4, 5, 6 and 7. *Anagyrus lapachosus* females lived on average 5 ± 3 days (range 2-12 days), and emerged with 57 ± 2 mature eggs (*e*). After emerging, a female produced 11 ± 1 eggs (*h*_0_) and the daily egg production rate was 1.330 ± 0.024 (*g*). Eggs that were not used on day *t*, were resorbed the next day in a proportion of 0.935 ± 0.028 (*r*). In the case of equations 5 and 7, for resorption to exist, the number of remaining eggs from one day to another had to exceed a threshold of 15 ± 1 eggs (*u*). Finally, just for equations 6 and 7, females presented a maximum egg storage capacity of 59 ± 2 eggs (*C*). The models *D*4, *D*5, *D*6, and *D*7 are detailed in S3 Table and represented in S5-S8 Figs.

**Fig 4.**
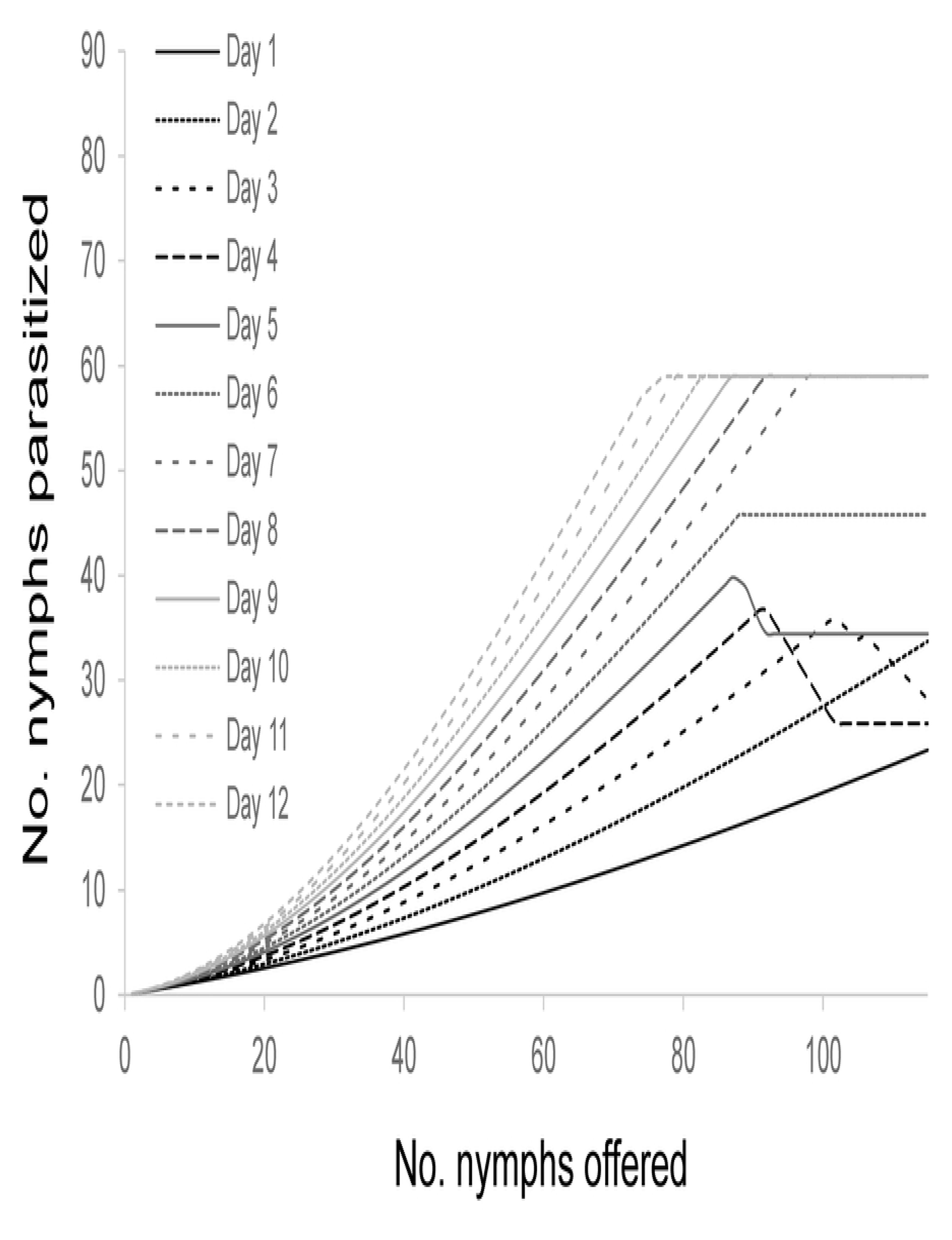
Estimated type III functional response of *Anagyrus lapachosus* females at different ages (days) of their lifespan (1-12 days) considering limited egg production and female experience. In equation *D* of the functional response module, the female gains experience throughout her life by interacting with the hosts, this is reflected in a daily increase in her attack rate.

## Discussion

We proposed a series of latent-variables Markovian models that comprised an integrated approach of functional response and egg production models to study the realized lifetime reproductive success of adult parasitoids. With this approach, insight was gained about the reproductive biology dynamics of two parasitoid species being considered as biological control agents of a cactus mealybug pest. The number of hosts parasitized by *A. cachamai* decreased with female age and depended on the number of mature eggs that were available for oviposition (Fig 3A and 3B), while host parasitism by *A. lapachosus* increased with female age and was modulated by its daily egg load and previous experience (Fig 4).

According to Vinson [60], parasitoid females show distinct oviposition behaviors, consisting of host location and evaluation, ovipositor insertion, host acceptance, oviposition, and host marking (chemical or mechanical to avoid superparasitism). Although host finding and attack cycle are inborn, experience gained during the oviposition process may result in an improvement in the skill of the females to locate and parasitize their host [61]. However, a parasitoids’ “motivation” is another element that is important in describing a parasitoids’ oviposition behavior, and may be influenced by factors such as the onset of hunger, egg load, presence of competitors and predators, as well as environmental changes [60]. This concept is assigned to the category of hidden or latent variables, which cannot be measured directly but only by its correlation with observable behavior [62]. With the models we developed, we explained the oviposition behavior of *A. cachamai* and *A. lapachosus* against variations in host densities in more detail than with the commonly used classical functional response models [1,63. Furthermore, by studying both species throughout their adult lives, we were able to analyze how age and previous experience with the host influenced their reproductive success.

Both species *A. cachamai* and *A. lapachosus* were determined to have a type III functional response. Holling [64] suggested that type III functional responses could be a consequence of parasitoid learning, however, his formulations of this behavior were not permanent. At low host densities, Holling’s model assumed that the contact of the parasitoid with the host would be so rare that the parasitoid could not develop a “search image” for the host. If host density increased, the frequency of contacts would rise and the parasitoid could become more responsive to the specific stimuli of the host. However, if the parasitoid does not encounter the host for a prolonged period of time, everything learned will be forgotten. The increased foraging behavior exhibited by females after parasitizing the first host may be due to a process known as associative learning [60]. Associative learning is identified as a females’ perception of chemical traces (semiochemicals) and/or physical stimuli of the host (visual or mechanical) after a full oviposition experience, and the parasitoid’s subsequent ability to find, recognize and accept (or reject) other hosts [65]. When a female is rewarded after a full oviposition experience, it learns that its foraging behavior in response to certain plant odors or host cues leads to finding a suitable host. Females of *A. cachamai* were “fast learners” after a single oviposition experience, although their response faded at 24 hours (Fig 3A and 3B). In contrast, females of *A. lapachosus* were “slow learners”, but they developed a long-term memory, since they showed an increase in their daily attack rate (Fig 4). Learning abilities and memory retention vary among parasitoid species and comes at a physiological cost [66,67. The different learning skills observed between *A. cachamai* and *A. lapachosus* females may be the result of their dissimilar reproductive strategies [67].

Thanks to the use of Markovian models combined with Bayesian statistics, it was possible to make an accurate description of the ovigeny strategy of *A. cachamai* and *A. lapachosus*. The number of mature eggs after emergence, egg production on the first day, egg production rate, the proportion of eggs resorbed, egg resorption threshold, and egg storage capacity was estimated for both species. The selected models confirmed that *A. cachamai* and *A. lapachosus* are synovigenic, coinciding with the results obtained by Aguirre et al. [42]. Our models indicated that *A. cachamai* females emerged with 56 ± 2 mature eggs and that their storage capacity was 58 ± 2 eggs, and that *A. lapachosus* females emerged with 57 ± 2 eggs and their storage capacity was 59 ± 2 eggs. Therefore, both species emerged with their maximum egg storage capacity.

Synovigenic parasitoids possess a variety of adaptations that reduce the risk of egg limitation and extend their lifespan (variable egg production rates, host acceptance or rejection, superparasitism of hosts, adjustable clutch size, egg resorption, host feeding) [25]. The egg production rate (*g*) of *A. cachamai* females decreased with increasing female age while the egg production rate for *A. lapachosus* increased with increasing female age. To the best of our knowledge, there are few studies that provide information about how egg production is affected by female age. In addition, most of the available information is for experimental designs where females received an excess of hosts. For example, Manzano et al. [68] reported that the egg production rate of *Cosmocomoidea annulicornis* (Ogloblin) (Hymenoptera: Mymaridae) females is affected by age. The lowest egg load observed is when the females are 1 and 12 hours old and the highest when females are 4, 5, and 8 hours old. [69]. On the other hand, in the egg parasitoid *Anagrus virlai* Triapitsyn (Mymaridae), the number of parasitized eggs decreases as females age, and wasps experience a double egg maturation process [30]. Palottini [62], using a similar experimental design and statistical analysis to the one we employed, found that the egg production rate of *Gonatocerus* sp. “clado 1” (Mymaridae) aff. *tuberculifemur* is 0.78, meaning that the egg production rate decreases with female age. We also determined that both species needed time to replenish their egg supply when the oviposition rate was high (S1-S8 Figs). *Anagyrus lapachosus* females had a lower egg resorption threshold (*u*) than *A. cachamai* females but shared the same egg storage capacity (*C*). Egg resorption by parasitoids may be a mechanism to remove unviable eggs [70] or to recycle nutrients [71]. Most likely, *A. cachamai* and *A. lapachosus* females experience egg resorption when host densities are too low or unsuitable to provide adequate oviposition opportunities.

Our data provide evidence that the risk of egg limitation was higher for *A. cachamai* females than *A. lapachosus*, since egg maturation declined with *A. cachamai* female age. *Anagyrus lapachosus* females presented two biological traits that gave them “flexibility” over *A. cachamai* females during the oviposition process: 1) increased egg production rate (*g*) with increasing female age; 2) female gain in experience over the course of her life when interacting with the host.

Functional response experiments are usually carried out for a short amount of time (1- 48 hours), ignoring that the wasp presents non-foraging behaviors until it is ready to begin host foraging (e.g. maturing or resorbing eggs, resting, grooming, exploring the experimental arena, etc.) [5]. The problem of non-foraging behaviors during functional response experiments can be addressed with the explicit inclusion of non-foraging mechanisms into the functional response models. Likewise, experimental trials should be of sufficient duration so that egg production, resting, and other normal non-foraging behaviors are expressed during the trial. This approach has the potential to fully address the problem arising from the expression of non-foraging behavior by parasitoids during functional response experiments, but it may result in complicated models that are challenging to fit to data and are difficult to interpret. In this work, thanks to the use of Markovian models combined with Bayesian statistics, it was possible to deal with non-foraging behavior when measuring a parasitoids’ functional response.

## Conclusions

The presented methodology has broad application and the potential to increase understanding of the reproductive biology of a wide variety of parasitoid species. From an applied perspective, our developed models have implications for the use of parasitoids as biological control agents. Unlike classical functional response methodology, we assessed candidate species according to their physiology and prior experiences. Using this methodological approach to predict the success of parasitoids as control agents will increase the amount of information obtained from the studied potential biological control species leading to more effective and safe agent selection.

## Acknowledgments

We thank Arabella Peard for reviewing a draft of the manuscript. Aguirre María Belén was the recipient of a PhD awarded by CONICET (Consejo Nacional de Investigaciones Científicas y Técnicas). Bruzzone Octavio is a research member of CONICET. This study was funded by the USDA-APHIS Farm Bill 19-8130-0852-IA (2020), and USDA-APHIS Invasive Species Coordination Program from 2014 to 2016, APH-HQ-16-0181.

## Supporting information

**S1 Fig.**
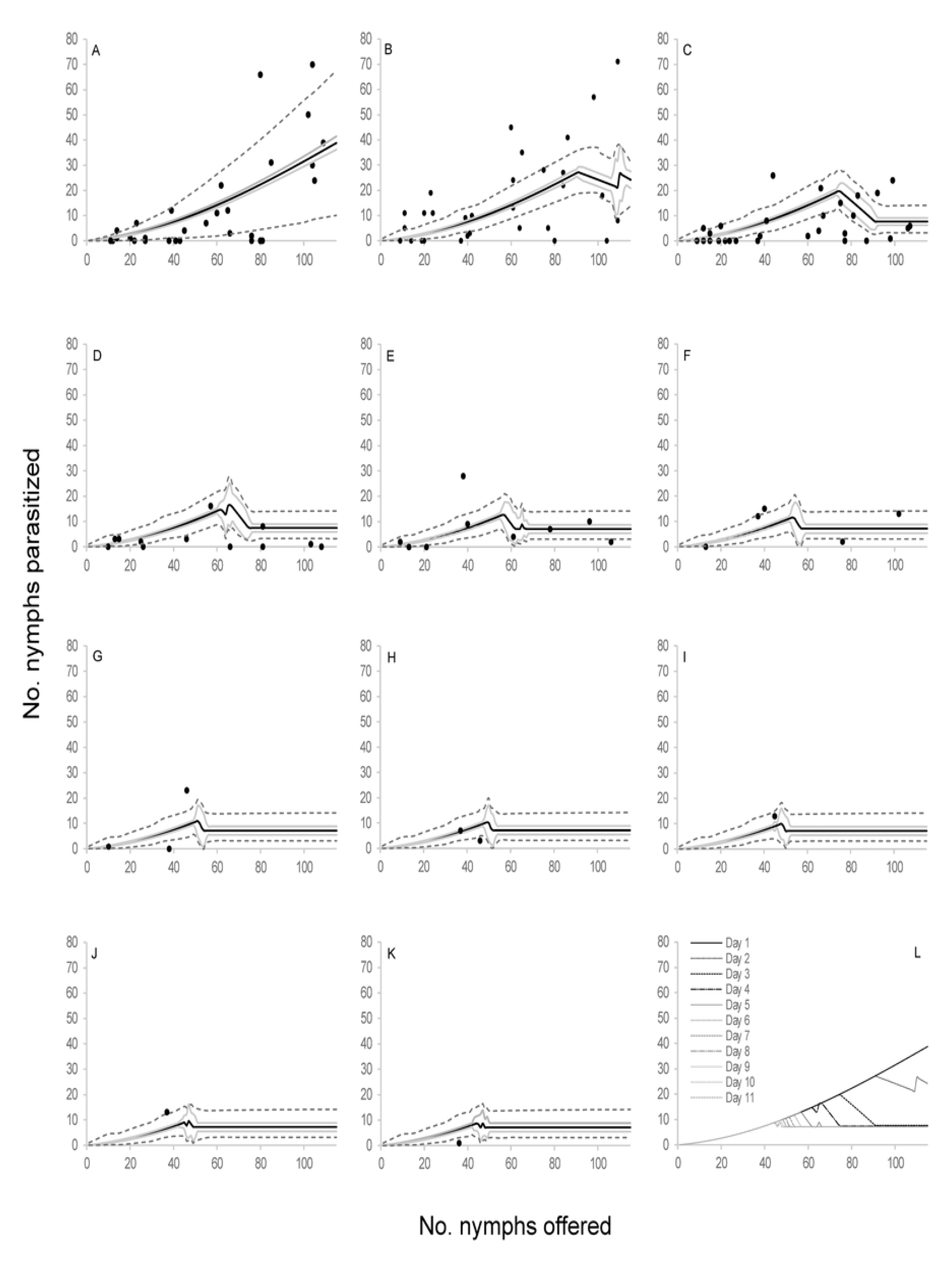
Observed functional response of the parasitoid *Anagyrus cachamai* attacking *Hypogeococcus* sp. nymphs. (A-K) Solid line indicates the mean estimation of functional response for model *C*5 at different ages of female lifespan (1-11 days), grey line indicates its credibility interval, and dashed line indicates the *a posteriori* credibility interval for individual measurements. Dark circles are the observed number of emerged parasitoids; (L) estimated functional response for model *C*5 from day 1 to11.

**S2 Fig.**
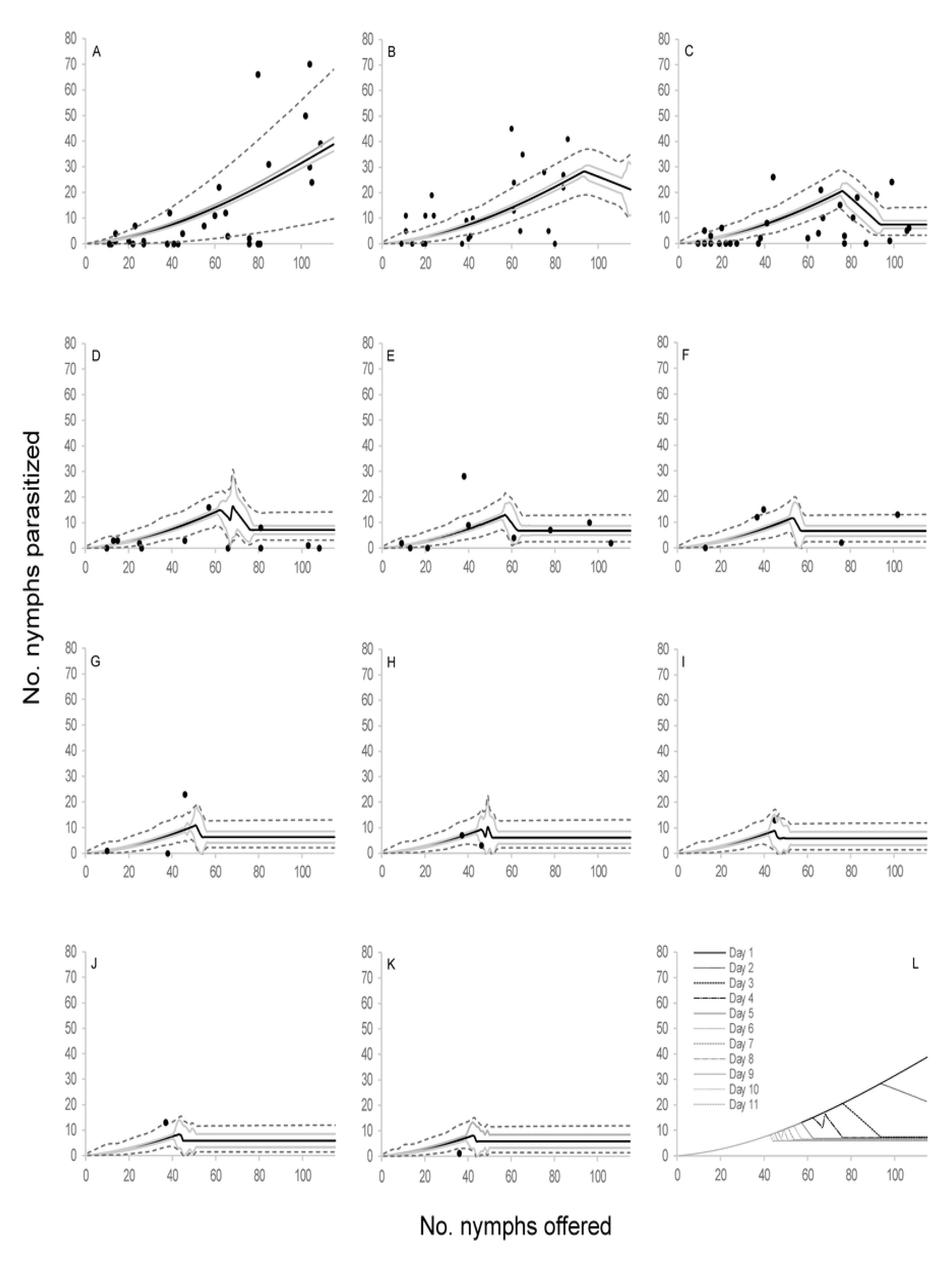
Observed functional response of the parasitoid *Anagyrus cachamai* attacking *Hypogeococcus* sp. nymphs. (A-K) Solid line indicates the mean estimation of functional response for model *C*7 at different ages of female lifespan (1-11 days), grey line indicates its credibility interval, and dashed line indicates the *a posteriori* credibility interval for individual measurements. Dark circles are the observed number of emerged parasitoids; (L) estimated functional response for model *C*7 from day 1 to11.

**S3 Fig.**
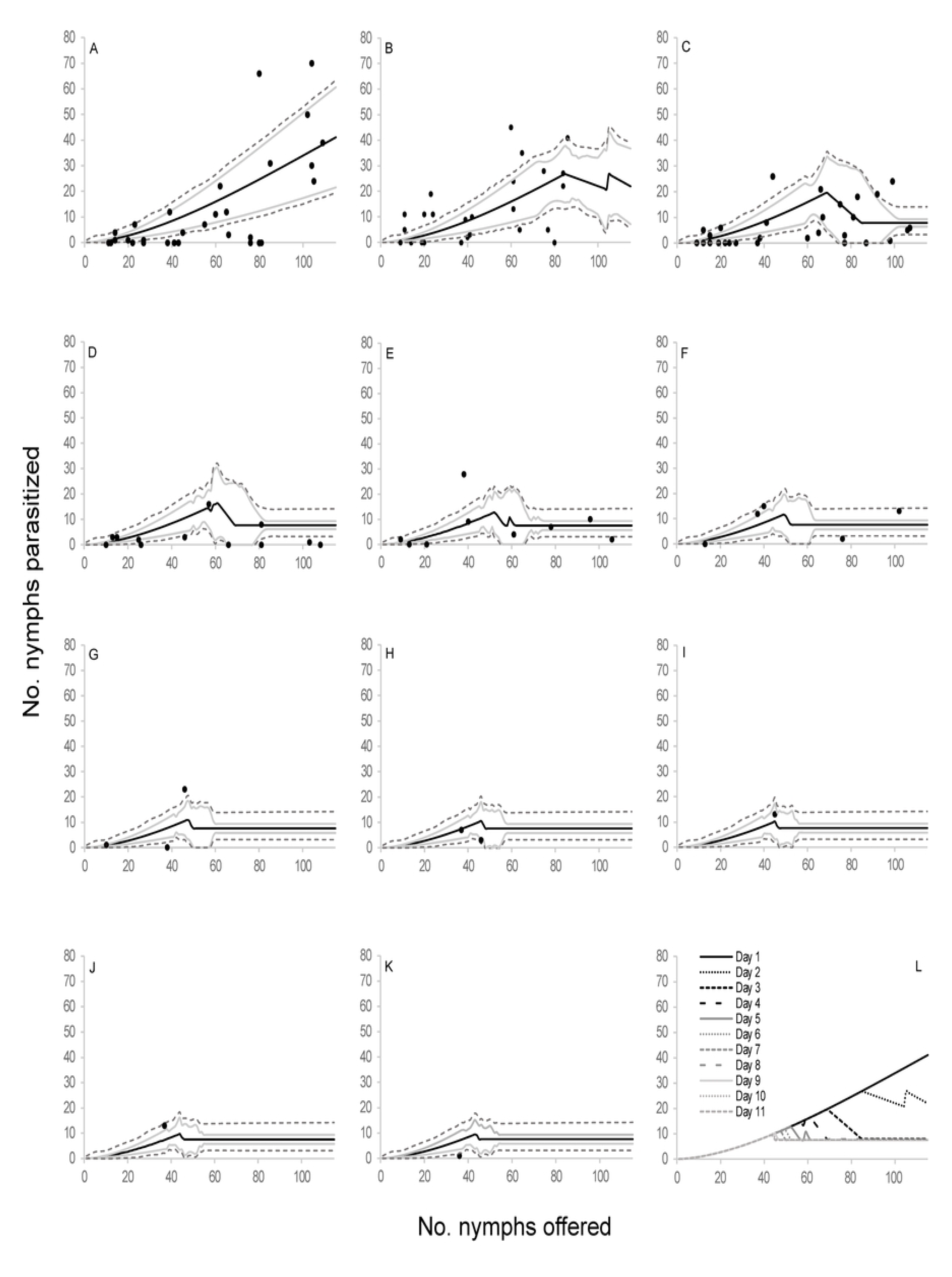
Observed functional response of the parasitoid *Anagyrus cachamai* attacking *Hypogeococcus* sp. nymphs. (A-K) Solid line indicates the mean estimation of functional response for model *E5* at different ages of female lifespan (1-11 days), grey line indicates its credibility interval, and dashed line indicates the *a posteriori* credibility interval for individual measurements. Dark circles are the observed number of emerged parasitoids; (L) estimated functional response for model *E*5 from day 1 to11.

**S4 Fig.**
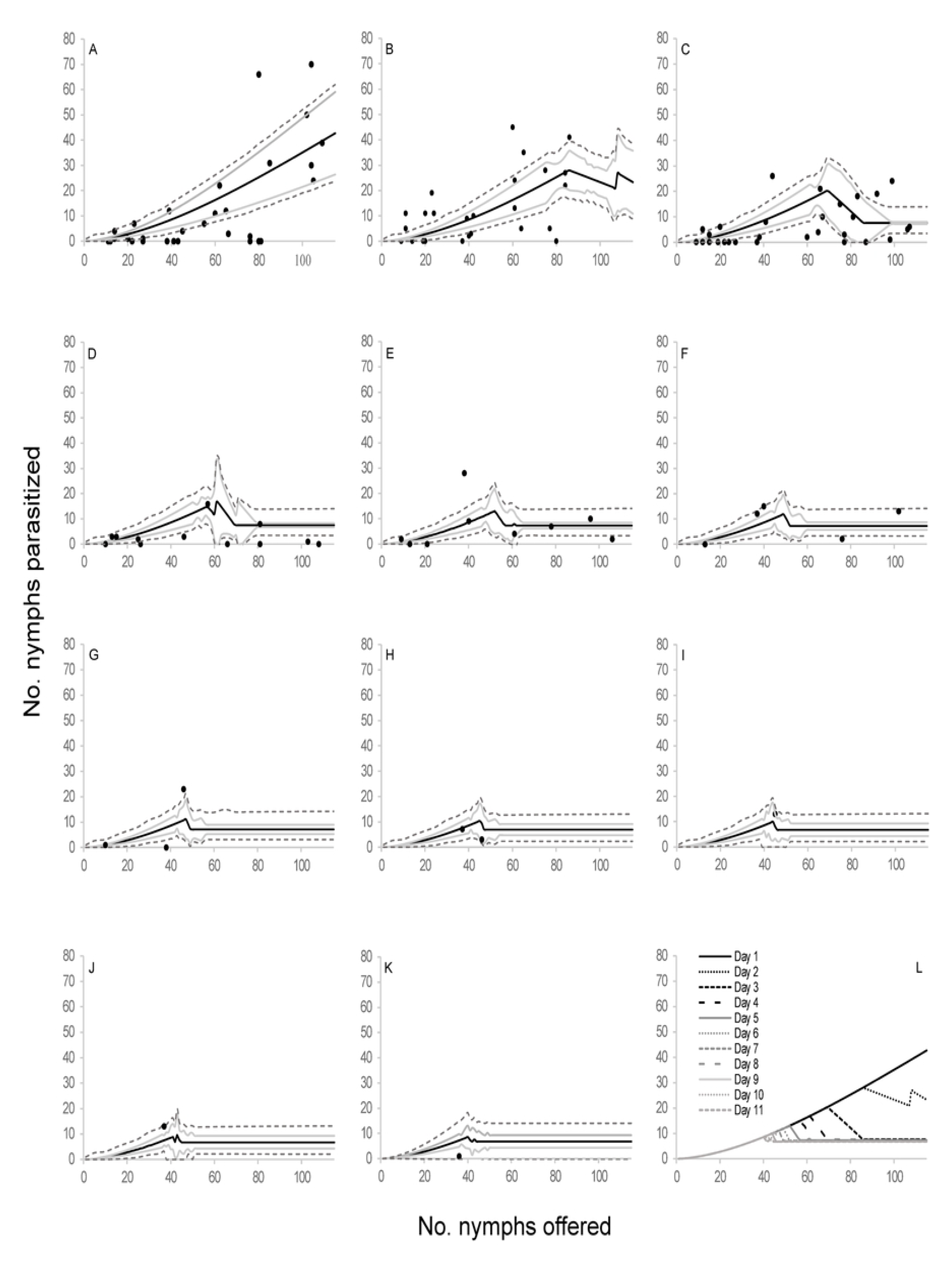
Observed functional response of the parasitoid *Anagyrus cachamai* attacking *Hypogeococcus* sp. nymphs. (A-K) Solid line indicates the mean estimation of functional response for model *E7* at different ages of female lifespan (1-11 days), grey line indicates its credibility interval, and dashed line indicates the *a posteriori* credibility interval for individual measurements. Dark circles are the observed number of emerged parasitoids; (L) estimated functional response for model *E*7 from day 1 to11.

**S5 Fig.**
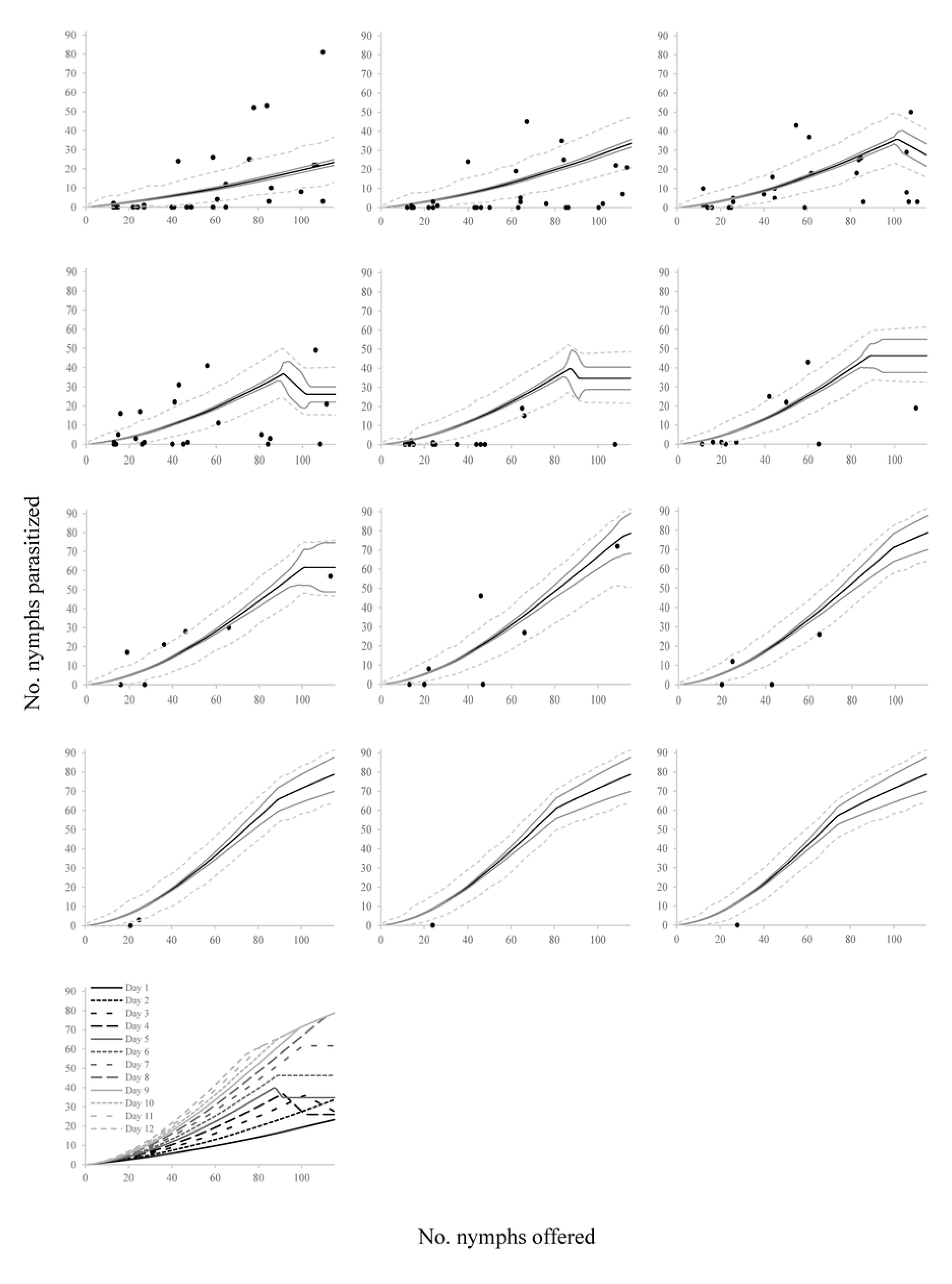
Observed functional response of the parasitoid *Anagyrus lapachosus* attacking *Hypogeococcus* sp. nymphs. (A-L) Solid line indicates the mean estimation of functional response for model *D*4 at different ages of female lifespan (1-12 days), grey line indicates its credibility interval, and dashed line indicates the *a posteriori* credibility interval for individual measurements. Dark circles are the observed number of emerged parasitoids; (M) estimated functional response for model *D*4 from day 1 to12.

**S6 Fig.**
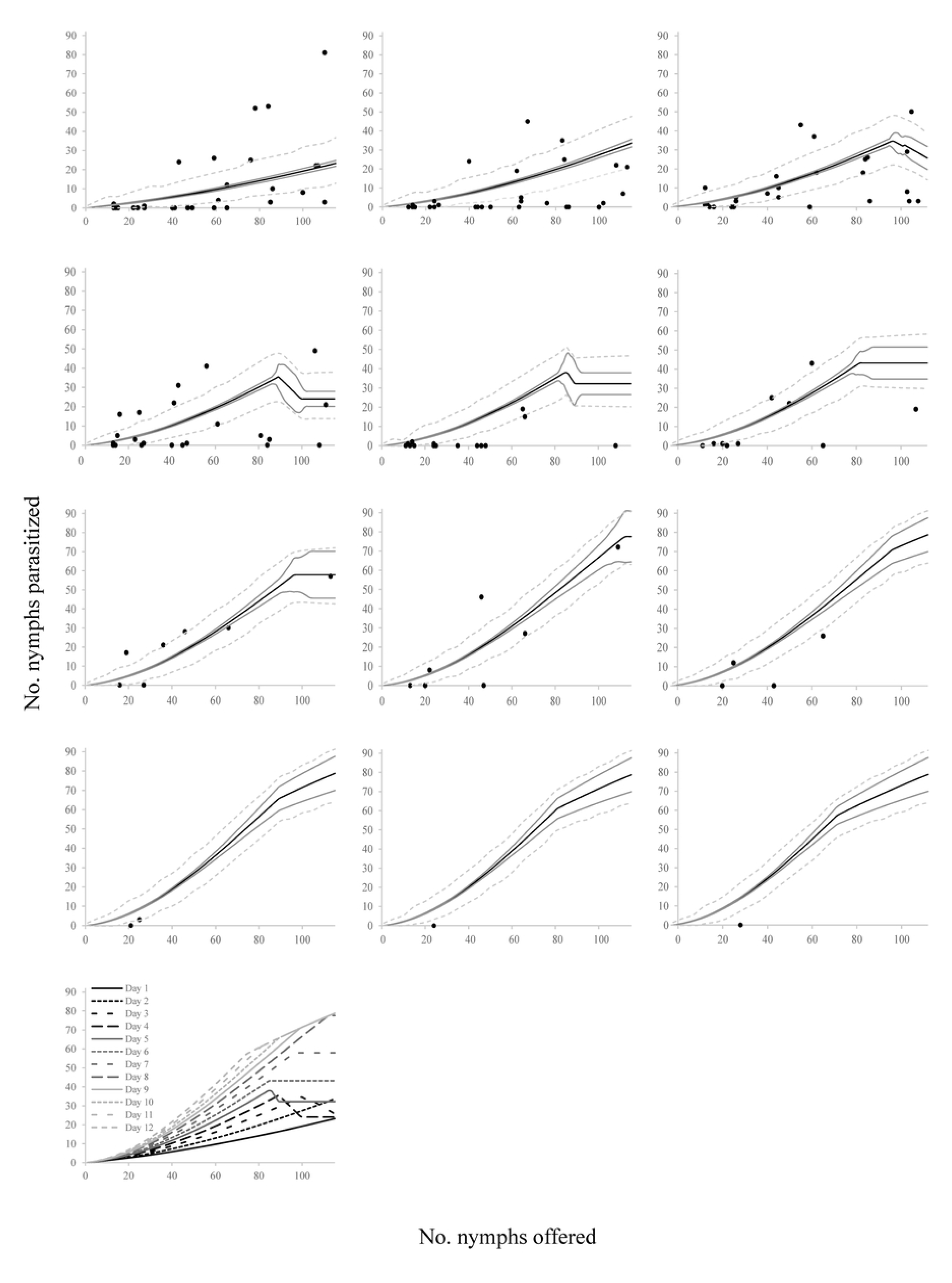
Observed functional response of the parasitoid *Anagyrus lapachosus* attacking *Hypogeococcus* sp. nymphs. (A-L) Solid line indicates the mean estimation of functional response for model *D*5 at different ages of female lifespan (1-12 days), grey line indicates its credibility interval, and dashed line indicates the *a posteriori* credibility interval for individual measurements. Dark circles are the observed number of emerged parasitoids; (M) estimated functional response for model *D*5 from day 1 to12.

**S7 Fig.**
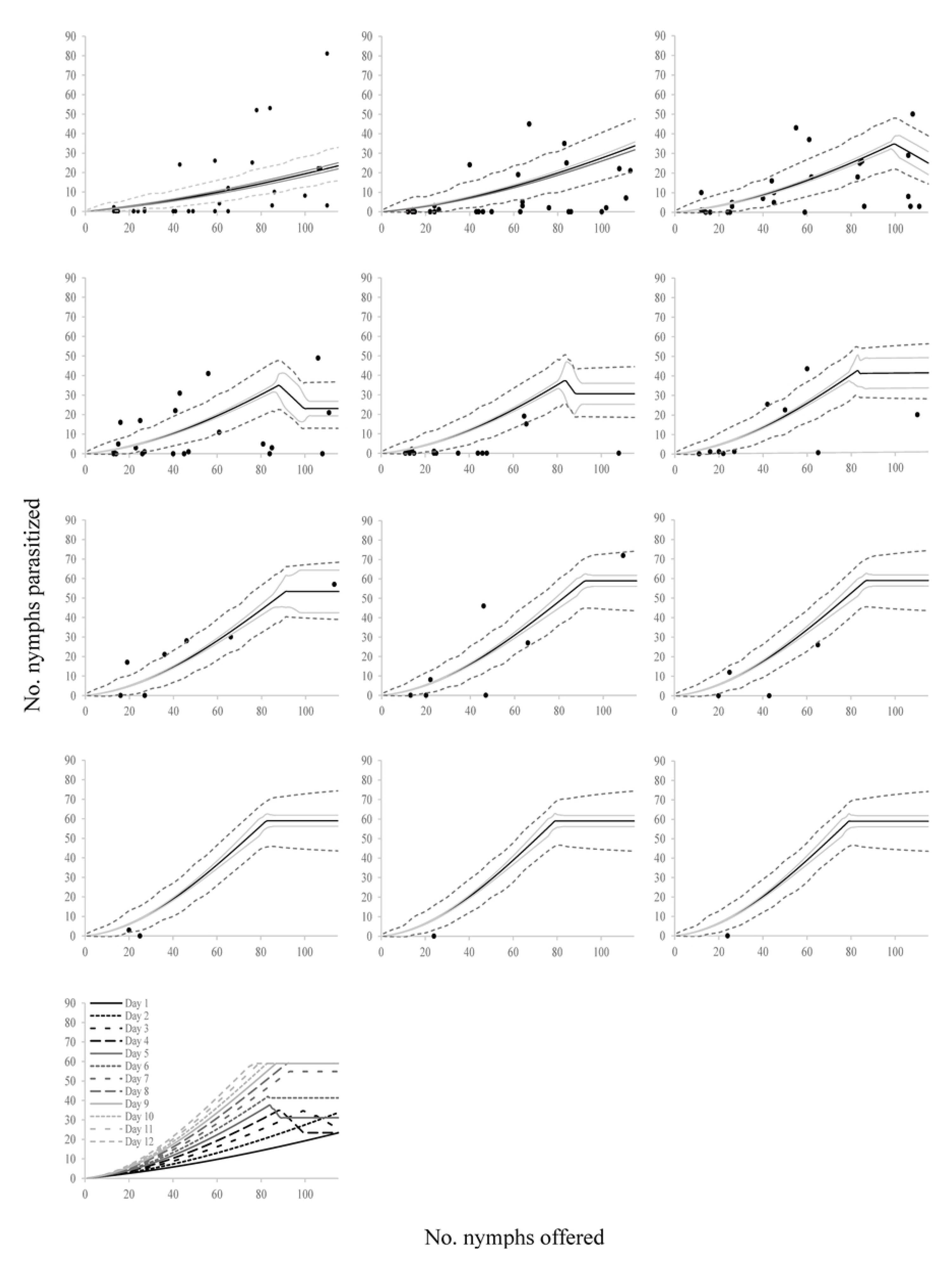
Observed functional response of the parasitoid *Anagyrus lapachosus* attacking *Hypogeococcus* sp. nymphs. (A-L) Solid line indicates the mean estimation of functional response for model *D*6 at different ages of female lifespan (1-12 days), grey line indicates its credibility interval, and dashed line indicates the *a posteriori* credibility interval for individual measurements. Dark circles are the observed number of emerged parasitoids; (M) estimated functional response for model *D*6 from day 1 to12.

**S8 Fig.**
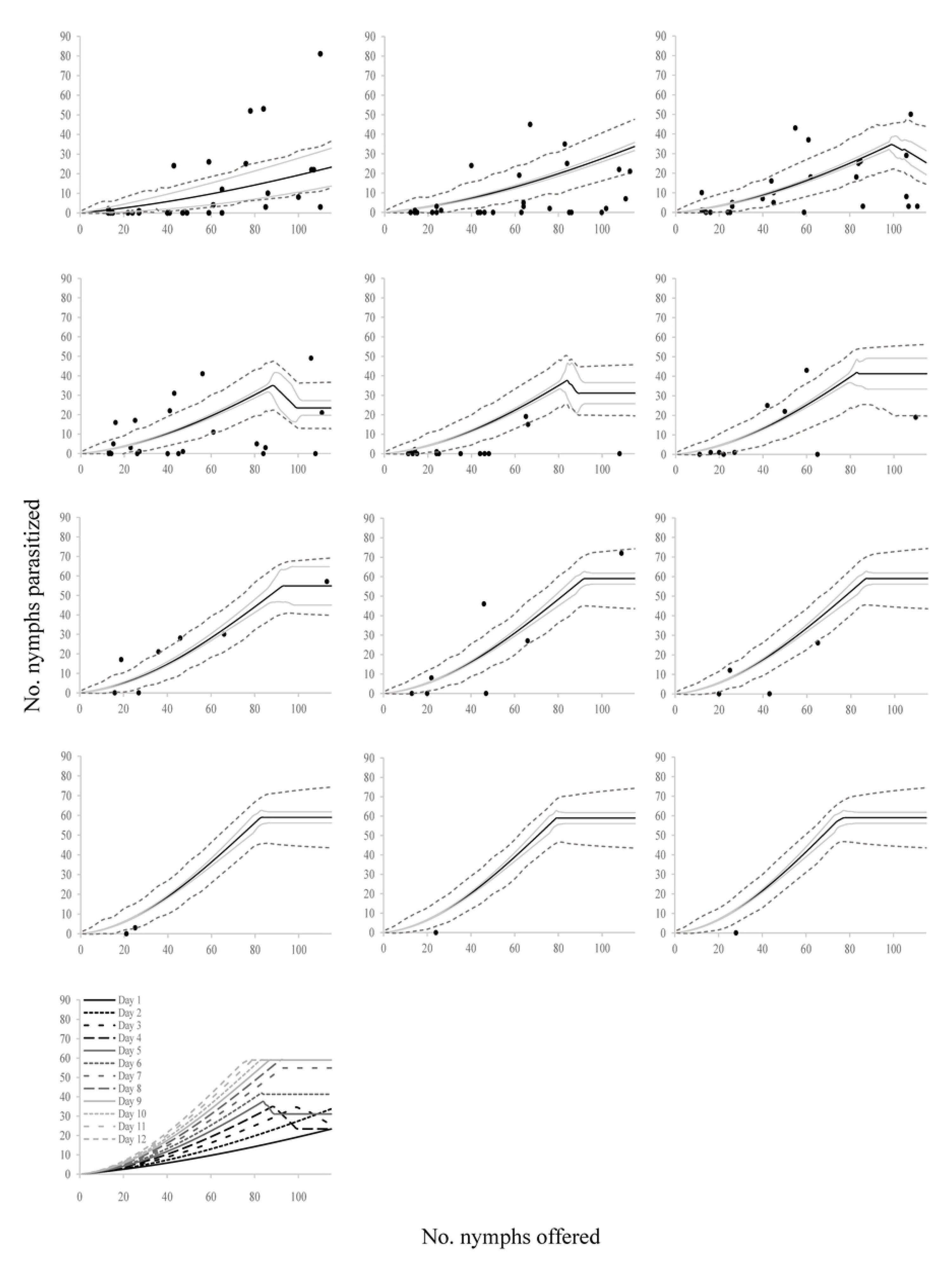
Observed functional response of the parasitoid *Anagyrus lapachosus* attacking *Hypogeococcus* sp. nymphs. (A-L) Solid line indicates the mean estimation of functional response for model *D*7 at different ages of female lifespan (1-12 days), grey line indicates its credibility interval, and dashed line indicates the *a posteriori* credibility interval for individual measurements. Dark circles are the observed number of emerged parasitoids; (M) estimated functional response for model *D*7 from day 1 to12.

**S1 Table. Deviance information criterion (DIC) of the 48 tested models.**

**S2 Table. Parameters of the selected models for the parasitoid species *Anagyrus cachamai*.**

**S3 Table. Parameters of the selected models for the parasitoid species *Anagyrus lapachosus*.**

**S1 File. Appendix.**

## Author Contributions

**Conceptualization:** María Aguirre, Guillermo Logarzo, Octavio Bruzzone.

**Data curation:** María Aguirre, Guillermo Logarzo, Serguei Triapitsyn, Octavio Bruzzone.

**Formal analysis:** María Aguirre, Guillermo Logarzo, Octavio Bruzzone.

**Investigation:** María Aguirre, Guillermo Logarzo, Octavio Bruzzone.

**Methodology:** María Aguirre, Guillermo Logarzo, Octavio Bruzzone.

**Project administration:** Guillermo Logarzo, Hilda Diaz-Soltero, Stephen Hight.

**Resources:** Hilda Diaz-Soltero, Stephen Hight.

**Software:** Octavio Bruzzone.

**Supervision:** Guillermo Logarzo, Octavio Bruzzone.

**Validation:** María Aguirre, Guillermo Logarzo, Octavio Bruzzone.

**Visualization:** María Aguirre, Guillermo Logarzo, Serguei Triapitsyn, Hilda Diaz-Soltero, Stephen Hight, Octavio Bruzzone.

**Writing – Original Draft Preparation**: María Aguirre, Guillermo Logarzo, Serguei Triapitsyn, Hilda Diaz-Soltero, Stephen Hight, Octavio Bruzzone.

**Writing – Review & Editing:** María Aguirre, Guillermo Logarzo, Serguei Triapitsyn, Hilda Diaz-Soltero, Stephen Hight, Octavio Bruzzone.

## Notes

### Competing Interest Statement

The authors have declared no competing interest.

